# Accumulation of *ph1* (*zip4-5B*) and *ph2* (*msh7-3D*) mutations fails to boost homoeologous recombination in hexaploid wheat

**DOI:** 10.1101/2025.10.27.684806

**Authors:** Camille Haquet, Isabelle Nadaud, Azahara Martin, Maria-Dolores Rey, Asma Ben Bouslah, Carus John-Bejai, Graham Moore, Pierre Sourdille

## Abstract

Diversification of the hexaploid (bread) wheat genetic pool using wild genetic resources relies on effective meiotic recombination (crossover) between wheat chromosomes and their counterparts from related species (homoeologues). However, crossover between homoeologues is normally suppressed by two major genes, *ZIP4-5B* (*Ph1*) and *MSH7-3D* (*Ph2*). We investigated the effect of introducing *zip4-5B* and/or *msh7-3D* mutations into interspecific hybrids derived from crosses between wheat and *Aegilops variabilis*. Single and double mutants were exploited in Chinese Spring (CS) and Cadenza (Cad) genetic backgrounds, as well as in a CS/Cad recombinant background. The number of univalents, bivalents and multivalents was scored from meiotic cells at metaphase I, from which numbers of chiasmata were deduced. We demonstrated a non-cumulative effect of simultaneous *zip4-5B* and *msh7-3D* mutations on homoeologous recombination, as homoeologous crossovers reached a maximum when *ZIP4-5B* alone was mutated. We also showed that hybrids carrying both the *zip4-5B* and *msh7-3D* mutations in the same genetic background exhibited more effective recombination compared to a double mutant in the CS/Cad recombinant background. The progression of meiosis was also monitored in the different interspecific hybrids mutants, revealing clear disruptions. Thus, our study significantly contributes to the optimization of the introgression of beneficial alleles from wild relatives into elite wheat germplasm; first by demonstrating the efficiency of *ZIP4-5B* and *MSH7-3D* mutations independently and in combination and second by elucidating the influence of the genetic background in which these mutations are present in an interspecific hybrid context.

## 1 Introduction

Bread wheat (*Triticum aestivum* L.) accounts for 20% of our global calorie consumption (Reynolds et al., 2012), yet concerns have been raised that the current rate of yield improvement may be insufficient to meet future demand, with projections indicating that at least a 40% increase will be required by 2050 (Charmet, 2011; Le Gouis et al., 2020; Ray et al., 2013).

Presently, our capacity to accelerate the rate of genetic improvement in wheat is constrained by (1) changes in agricultural practices for economic or regulatory reasons, such as the need to reduce mineral nitrogen fertilizer application, or measures to improve agricultural sustainability such as reductions in pesticide usage (Peltonen-Sainio et al., 2015); (2) climate change, which is becoming increasingly important and threatens yield potential as well as the stability of yield (Brisson et al., 2010; Le Gouis et al., 2020; Porter and Semenov, 2005; Tester and Langridge, 2010); (3) slowing of genetic gains due to environmental issues such as drought stress (Manès et al., 2012; Touzy et al., 2019). This suggests that new wheat varieties must be more tolerant to environmental stresses, especially drought and heat, and less dependent on fertilizers, insecticides or fungicides.

One way to face this challenge is to tap into wheat wild relatives, which serve as a large reservoir of unexploited alleles/genes, the value of which has been demonstrated in the genetic improvement of many agronomically important traits (Feuillet et al., 2008; King et al., 2024; Marcussen et al., 2014). Actually, it has been demonstrated that the introgression of genetic material from wild relatives has occurred during the evolution of wheat without the intervention of man (Cheng et al., 2019). From the 1970s onward, there has been a concerted effort by wheat researchers to hybridize such wild species with domesticated wheat with the aim of introducing novel beneficial diversity (Laugerotte et al., 2022). This resulted in a reshuffling of the wheat genome, and produced numerous structural variants larger than 5 Mb in most modern cultivars (Balfourier et al., 2019). Some of these wild relative introgressions have been widely used in wheat breeding programs like the 1RS/1BL wheat/rye (*Secale cereale*) translocation (Cai and Liu, 1989) and the 2A/2N introgression derived from *Aegilops ventricosa* (Helguera et al., 2003).

The main challenge in wheat wild relative introgression remains the exchange between wheat and wild relative chromosomes. Such exchanges rely on meiotic recombination, a process common to all sexually reproducing eukaryotes, and mandatory for viable and fertile haploid gamete production (for review, see Mercier et al., 2015). Meiotic recombination, also called crossover (CO) is tightly controlled, both in the number and location of recombination events (reviewed in Girard et al., 2023). Moreover, because bread wheat is an allopolyploid species (genome AABBDD; 2n = 6x = 42), deriving from two natural interspecific hybridizations of three closely related diploid species (*T. urartu* (AA), *Ae. speltoides* (SS related to BB), and *T. tauschii* (DD); Marcussen et al., 2014), there is an additional layer of control preventing COs between related homoeologous chromosomes within the wheat genome (Mason and Wendel, 2020; Sourdille et al., 2025).

In bread wheat, early genetic and cytogenetic studies revealed the presence of at least two loci, *Ph1* and *Ph2*, located on chromosome-arms 5BL and 3DS respectively, controlling recombination between its homoeologues (Mello-Sampayo, 1971; Mello-Sampayo and Lorente, 1968; Riley and Chapman, 1958; Sears and Okamoto, 1958). A large deletion encompassing *Ph1* was developed by Sears in Chinese Spring (CSph1b, 1977), covering nearly 60Mb and ∼1200 genes (Martín et al., 2018). However, the line carrying this deletion has accumulated extensive rearrangements over the years, including additional deletions, chromosome exchanges and duplications (Martín et al., 2018), thereby reducing the efficiency of the introgression process during breeding. Previous studies delimited the *Ph1* locus to a region containing a cluster of *CDK2-like* genes, as well as a block of heterochromatin duplicated from chromosome 3B (Al-Kaff et al., 2008; Griffiths et al., 2006; Martín et al., 2017). This heterochromatin block contained a copy of ZIP4, a protein required for the formation of type-I COs (Chelysheva et al., 2007). More recently, exploitation of Ethyl-Methyl Sulfonate (EMS), as well as CRISPR-Cas9 mutants of this additional-copy of ZIP4 on 5B (*zip4-5B*, originally named *TaZip4-B2*; Rey et al., 2017), showed that *zip4-5B* mutants mimic the original *ph1* Sears’ phenotype, exhibiting a significantly increased number of homoeologous COs (Alabdullah et al., 2021; Martín et al., 2021; Rey et al., 2018, 2017). These mutants revealed that this additional *ZIP4-5B* copy is also required for complete synapsis of homologues, with loss of this copy leading to incomplete synapsis (Draeger et al., 2023). Although the exact mode of action of *ZIP4-5B* remains unclear, evidence from tetraploid and hexaploid wheat suggests that it promotes early homologous synapsis, thereby reducing the opportunity for homoeologues to synapse and then crossover later in meiosis (Draeger et al., 2023; Martín et al., 2014). Interestingly, it was shown that different types of *zip4-5B* mutants exhibit distinct meiotic behaviours, with some retaining a high level of fertility despite facilitating homoeologous recombination, demonstrating a dual function for *ZIP4-5B* (Martín et al., 2021, 2014).

As regards to *Ph2*, two mutants were initially produced in the reference cultivar Chinese Spring: an irradiation mutant (*ph2a*; Sutton et al., 2003), with a deletion covering ∼125 Mb of the distal part of the short arm of chromosome 3D (Svačina et al., 2020), and an EMS mutant (*ph2b*; Wall et al., 1971). Previous attempts to clone *Ph2* suggested several different genes as potential candidates, including WM1 (Ji and Langridge, 1994; Whitford, 2002), WM3 (Letarte, 1996), WM5 (Dong et al., 2005) and *TaMsh7* (Dong et al., 2002; Lloyd et al., 2007). Recently, *MSH7-3D* (originally named *TaMSH7-D1*) was found to correspond to *Ph2* (Serra et al., 2021), coding for a plant specific MSH7 protein (Mut-S homolog 7) belonging to the DNA Mismatch Repair (MMR) complex (Culligan and Hays, 2000). Interestingly, MSH7 has also been found to be a key factor in homoeologous recombination in tomato (*Solanum lycopersicum*; Tam et al., 2011). It has been suggested that the *MSH7-3D* protein could play a crucial role during the early steps of recombination, by rejecting heteroduplex single-strand invasion between homoeologues (Serra et al., 2021).

Hybrids between wild relatives and wheat can be used to score homoeologous chiasmata (the visible manifestation of crossover events), as there are no homologous chromosomes present in such hybrids, only homoeologues. Previous studies utilized such hybrids to assess the interaction between *Ph1* and *Ph2* on homoeologous CO. Mello-Sampayo and Canas (1973) assessed homoeologous chiasmata in crosses involving aneuploid wheats, missing either full chromosomes (3A, 3D or 5B) or chromosome-arms (3AS, 3BS, 3DS and 3AS + 3DS), and *Aegilops sharonensis* (genome SshSsh) or rye (*Secale cereale*; genome RR). They observed that the number of homoeologous chiasmata was greatest when chromosome 5B was missing, compared to when either chromosome 3D or chromosome-arm 3DS was missing. In the absence of chromosome 5B, 50% of the 28 homoeologous chromosomes were linked by chiasmata on both their chromosome arms. The additional loss of 3DS did not increase chiasma formation; instead, it caused a slight reduction in the average number of chiasmata per cell (15.07 vs 13.84, respectively).

Ceoloni and Donini (1993) applied a similar approach, exploiting hybrids with *Aegilops variabilis* (UUSvSv; 2n = 28) or rye to evaluate meiotic chromosome configurations, with the aim of deducing the extent of homoeologous chiasmata. Contrary to Mello-Sampayo and Canas (1973), they observed an 8% higher level of homoeologous chiasmata when both *Ph1* and *Ph2* were absent, compared to when just *Ph1* was absent. This difference was due to an increase in chiasmata linking multiple chromosomes. However, they highlighted the difficulty of rapidly and unambiguously screening homozygous and heterozygous double *ph1/ph2* mutants, as meiosis-based checks were sometimes unreliable due to background effects from large deletions.

In this study we took advantage of the recent cloning of both *Ph1* and *Ph2* (Rey et al., 2017; Serra et al., 2021), to generate a large and comprehensive set of crosses involving these two loci and to design specific markers for each locus. Using these markers, we screened crosses between *zip4-5B* (*ph1*) and/or *msh7-3D* (*ph2*) mutants and *Ae. variabilis*, to identify specific single and double mutations. This approach enabled us to assess the individual and combined contributions of *ZIP4-5B* and *MSH7-3D* to homoeologous CO formation, focusing on their specific effects rather than the broader consequences of deletions or loss of whole chromosomes covering multiple genes.

## 2 Materials & Methods

### 2.1 Plant material

Wheat-*Aegilops variabilis* interspecific hybrids were produced using five different wheat mutants in two genetic backgrounds: Chinese Spring (CS) and Cadenza (Cad). For CS, *ph1b* (deletion of 59.3 Mb; (Sears, 1977) and *ph2b* (EMS mutant; Wall et al., 1971) reference mutants were used. For Cad, three EMS mutants from the John Innes Centre collection (https://www.jic.ac.uk/research-impact/germplasm-resource-unit/; Krasileva et al., 2017) were used: two *zip4-5B* mutants (Cad0348 and Cad1691; (Martín et al., 2021; Rey et al., 2017) and one *msh7-3D* mutant (Cad2006; Serra et al., 2021). Wild-type accessions for CS and Cad were used as control.

The crossing strategy used to generate the mutant collection is illustrated in Figure 1. Within each genetic background (CS and Cad), single wheat mutants were first inter-crossed to generate *ph1b/ph2b* double mutants in CS or *zip4-5B/msh7-3D* double mutants in Cad (Fig. 1A a,b). These double mutants were then crossed between backgrounds to produce CS x Cad double mutants (Fig 1A c). Single *ph1* and *ph2* mutants from the two different genetic backgrounds were also crossed (Fig. 1A d,e). Unfortunately, the cross between the ph1 mutants was not successful, and we were therefore unable to obtain the CS *ph1b*/Cad *zip4-5B* genotype (Fig. 1A e). In parallel, the CS and Cad wildtypes were also crossed (Fig 1A f). The resulting hybrids (CS/Cad (f), CS *ph2b*/Cad *msh7-3D* (d), CS *ph1b* Cad *zip4-5B* / CS *ph2b* Cad *msh7-3D* (c)) were finally crossed with *Aegilops variabilis* (Aev; ERGE code 26248) to generate interspecific haploid hybrids (Fig. 1B). Twenty-nine interspecific haploid hybrids were obtained following this scheme and evaluated (Supplementary Table 1). Molecular analyses were conducted to check their mutant status: wildtype, CS *ph2b* or Cad *msh7-3D* single mutant; CS *ph1b/ph2b* or Cad *zip4-5B/msh7-3D* double mutant; or CS/Cad recombined double mutant (CS *ph1b*/Cad *msh7-3D* or Cad *zip4-5B*/CS *ph2b*). After analysis, the wild-type control individuals (Cad and CS x Aev) were not retained because the genotyping was not successful. Finally, 12 of the 29 interspecific hybrid haploid plants were randomly selected for further analyses (Supplementary Table 1). For Cad and CS controls and the single mutants *ph1* (CS *ph1b* and Cad *zip4-5B*) crossed with Aev, we used data from 21 hybrids obtained in previous experiments (Martin, Rey and Moore, unpublished results; Fig. 1C)).

**Figure 1.**
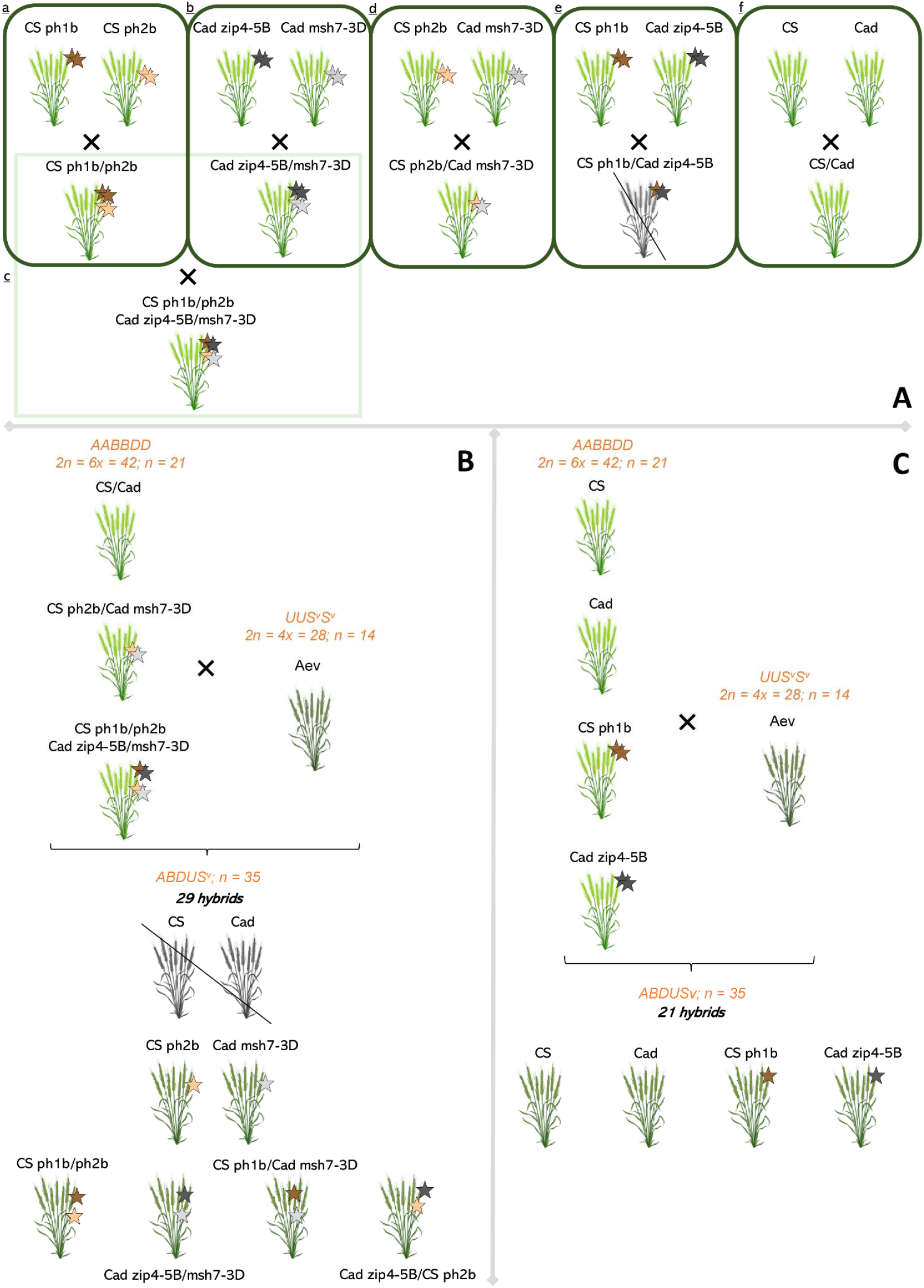
Crossing scheme performed between wheat lines (*Triticum aestivum*, 2n = 6x = 42) with or without mutations at the *Ph1* and/or *Ph2* locus and a related species, *Aegilops variabilis* (2n = 4x = 28) (A) Single mutants for the *ph1* and/or *ph2* loci were generated and inter-crossed to produce double mutants in the same genetic background for (a) CS and (b) Cad varieties. Double mutants were then crossed to produce CS × Cad double mutant (c). The CS and Cad *ph2* single mutants were crossed to obtain an individual carrying *ph2* from both backgrounds (d), same with *ph1* (e). The cross between the *ph1* (e) mutants was unsuccessful. The CS and Cad controls were also crossed (f). (B) Subsequently, plants derived from (c) (d) and (f) crosses were hybridised with *Aegilops variabilis* (Aev). The resulting 29 interspecific hybrids carry a genome of 35 chromosomes (ABDUSv). Unfortunately, the control interspecific hybrids CS and Cad were not retained for further analyses. (C) Since the controls CS and Cad interspecific hybrids, as well as the single ph1 mutants, were not available, we used individuals from a previous experiment. Wild-type CS and Cad, and single mutants for the *ph1* locus were crossed with *Aegilops variabilis* (Aev). The resulting 21 hybrids carry an allo-haploid genome of 35 chromosomes (ABDUSv).

In total, 33 interspecific haploid hybrids with the following genotypes at *Ph* loci were phenotypically evaluated (Supplementary Table 2):

▪ five wild-type Cad;
▪ two wild-type CS;
▪ twelve Cad *zip4-5B* single mutants;
▪ two CS *ph1b* single mutants;
▪ one Cad *msh7-3D* single mutant;
▪ one CS *ph2b* single mutant;
▪ two CS *ph1b/ph2b* double mutants;
▪ three Cad *zip4-5B/msh7-3D* double mutants;
▪ three recombined CS *ph1b*/Cad *msh7-3D* double mutants;
▪ two recombined Cad *zip4-5B*/CS *ph2b* double mutants.

Seeds were sown in potting soil and grown until the three-leaf stage. They were then transferred in a vernalization chamber at 6°C ± 1°C for two months under an 8 h light/16 h dark photoperiod. After vernalization, plants were transplanted into 4-litre pots containing Nutricote (Fertil, a commercial progressive release fertilizer), and placed in a greenhouse under a 16 h light/8 h dark photoperiod at 23°C/18°C (day/night). A small piece of leaf was collected for DNA extraction at the time of transplanting.

### 2.2 Molecular analyses

DNA was isolated from 100 mg of fresh leaves using a magnetic beads protocol (Sbeadex mini plant kit, LGC, Teddington, United Kingdom), with concentrations adjusted to 10ng/µL. PCR was performed in 11 µL (5 µL Master Mix AmpliTaq Gold™ 360, 1 µL Master Mix 360 GC Enhancer, 0.5 µL primers mix (forward and reverse at 10 µM, 1.5 µL water, 3 µL of DNA at 10ng/µL) under the following conditions: initial denaturation at 95°C for 10 min; 35 cycles of denaturation at 95°C for 30 s, annealing at Tm for 30 s, and elongation at 72°C for 50 s; followed by a final extension at 72°C for 10 min. Sequences of primers and Tm are given in Supplementary Table 3. The presence/absence of *Ph1* and/or *Ph2* mutations were scored on 1.2% agarose gel prepared in TAE 1X (Tris 0.04 M, Acetate 0.04 M, EDTA 0.001 M) and run during 30 minutes at 100V.

### 2.3 Cytogenetic studies and statistics

Anther extraction, meiocyte preparation and microscopy observations were as described in (Bazile et al., 2024). Briefly, tillers with immature inflorescences were collected in the morning and placed immediately on ice. Spikes were carefully removed from the sheath, and the three anthers of each flower extracted. One anther was stained with Acetocarmine (10 g/L Carmin, 45% acetic acid) and observed under a microscope to check for the meiotic stage. The remaining two synchronized anthers were fixed in Carnoy solution (EtOH 100% - glacial acetic acid; v/v 3:1) for 48 hours, followed by storage in EtOH 70% at 4°C. To analyse meiotic progression of different hybrids from zygotene to the tetrad stage in detail, one anther from each stage was placed on a poly-L-lysine-coated slide with a drop of acetocarmine and opened using two roll pins under binocular to expose the meiocytes. A drop of 45% acetic acid was applied to remove acetocarmine, and slides were immersed in liquid nitrogen, air-dried briefly, and mounted with Vectashield-DAPI (Eurobio-Ingen). Images were captured using an Axio Observer Z1 fluorescence microscope with Zen software (Carl Zeiss microscopy).

The number of cells examined per genotype ranged from 48 to 139 (Table 1). Numbers of univalents, bivalents (ring and rod) and multivalents (including complex pairing configurations) were counted manually, and numbers of chiasmata deduced from these counts, according to the method described in Supplementary Figure 1. For each dataset, normality of distribution was checked using the normality test of Shapiro and Wilk with default values. Since the data did not follow a normal distribution, a non-parametric Mann-Whitney test was applied.

**Table 1:** <u>Meiotic configurations in wheat–*Ae*. *variabilis* hybrids with different genotypes at the *Ph1* and/or *Ph2* loci in Chinese Spring (CS) or Cadenza (Cad) backgrounds. Values represent the mean ± standard deviation for the number of chiasmata at metaphase I: univalents, rod bivalents, ring bivalents and multivalents. n = the number of cells counted.</u>

| Genotype | Chiasmata | Univalents | Rod bivalents | Ring bivalents | Multivalents | n |
| --- | --- | --- | --- | --- | --- | --- |
| **Cad** | 3.25 ± 3.22 <sup>(a)</sup> | 30.23 ± 4.22 <sup>(g)</sup> | 1.53 ± 1.35 <sup>(a)</sup> | 0.86 ± 1.28 <sup>(a)</sup> | 0 ± 0 <sup>(a)</sup> | 80 |
| **CS** | 3 ± 2.46 <sup>(a)</sup> | 30 ± 3.65 <sup>(g)</sup> | 2 ± 1.59 <sup>(a)</sup> | 0.5 ± 0.98 <sup>(a)</sup> | 0 ± 0 <sup>(a)</sup> | 72 |
| **Cad *msh7-3D*** | 10.31 ± 3.07 <sup>(b)</sup> | 21.78 ± 3.55 <sup>(f)</sup> | 2.88 ± 1.41 <sup>(b)</sup> | 3.71 ± 1.6 <sup>(bcde)</sup> | 0.02 ± 0.14 <sup>(a)</sup> | 51 |
| **CS *ph2b*** | 13.88 ± 3.47 <sup>(c)</sup> | 15.94 ± 3.47 <sup>(e)</sup> | 4.73 ± 1.85 <sup>(c)</sup> | 4.08 ± 2.11 <sup>(de)</sup> | 0.48 ± 0.65 <sup>(bc)</sup> | 48 |
| **Cad *zip4-5B*** | 21.21 ± 3.33 <sup>(f)</sup> | 8.85 ± 3.47 <sup>(b)</sup> | 4.16 ± 1.93 <sup>(c)</sup> | 7.7 ± 2.3 <sup>(g)</sup> | 0.73 ± 0.87 <sup>(c)</sup> | 164 |
| **CS *ph1b*** | 19.4 ± 3.17 <sup>(e)</sup> | 9.69 ± 2.6 <sup>(c)</sup> | 4.78 ± 2.35 <sup>(c)</sup> | 5.85 ± 2.52 <sup>(f)</sup> | 1.25 ± 1.16 <sup>(d)</sup> | 67 |
| **Cad *zip4-5B/msh7-3D*** | 17.54 ± 2.92 <sup>(d)</sup> | 8.4 ± 3.8 <sup>(b)</sup> | 8.07 ± 2.38 <sup>(f)</sup> | 3.37 ± 1.72 <sup>(bc)</sup> | 1.14 ± 1.08 <sup>(d)</sup> | 139 |
| **CS *ph1b/ph2b*** | 19.63 ± 2.89 <sup>(e)</sup> | 6.63 ± 2.53 <sup>(a)</sup> | 7.68 ± 2.4 <sup>(ef)</sup> | 4.56 ± 2.16 <sup>(e)</sup> | 1.28 ± 1.32 <sup>(d)</sup> | 57 |
| **Cad *zip4-5B/CS ph2b*** | 14.32 ± 4.4 <sup>(c)</sup> | 13.29 ± 5.95 <sup>(d)</sup> | 7.03 ± 2.37 <sup>(de)</sup> | 3.11 ± 1.87 <sup>(b)</sup> | 0.46 ± 0.72 <sup>(b)</sup> | 98 |
| CS *ph1b*/Cad *msh7-3D* | 16.69 ± 3.29 <sup>(d)</sup> | 10.79 ± 4.82 <sup>(c)</sup> | 6.54 ± 2.33 <sup>(d)</sup> | 3.75 ± 1.71 <sup>(cd)</sup> | 1.13 ± 1.23 <sup>(d)</sup> | 138 |

## 3 Results

In total, 33 interspecific wheat-*Ae. variabilis* allo-haploid hybrids were generated, using wheat mutants in Chinese Spring (CS) or Cadenza (Cad) backgrounds affecting the *Ph1* and/or *Ph2* loci, together with their corresponding wild-type controls (see Materials and Methods; Supplementary Table 2). All plants showed normal development, with the exception of plant 13D (a CS *ph1b/ph2b* double mutant derived interspecific hybrid), which exhibited a slight delay in flowering compared to the others (Supplementary Figure 2). Three spikes from each hybrid were bagged to check for self-fertility. As expected, no seed was produced, confirming the sterility of the allo-haploid interspecific hybrids. Given the genomic constitution of bread wheat (AABBDD; 2n = 6x = 42; n = 21) and *Ae. variabilis* (UUSvSv; 2n = 4x = 28; n = 14), interspecific hybrid meiocytes must contain 35 chromosomes.

### 3.1 Analysis of interspecific hybrids with *ph1* or *ph2* mutations at meiotic metaphase I

Numbers of univalents, bivalents (ring and rod) and multivalents (more than two chromosomes linked by chiasmata) at metaphase I stage were scored for each interspecific hybrid, and total number of homoeologous chiasmata was subsequently deduced (Table 1; Figure 2).

**Figure 2.**
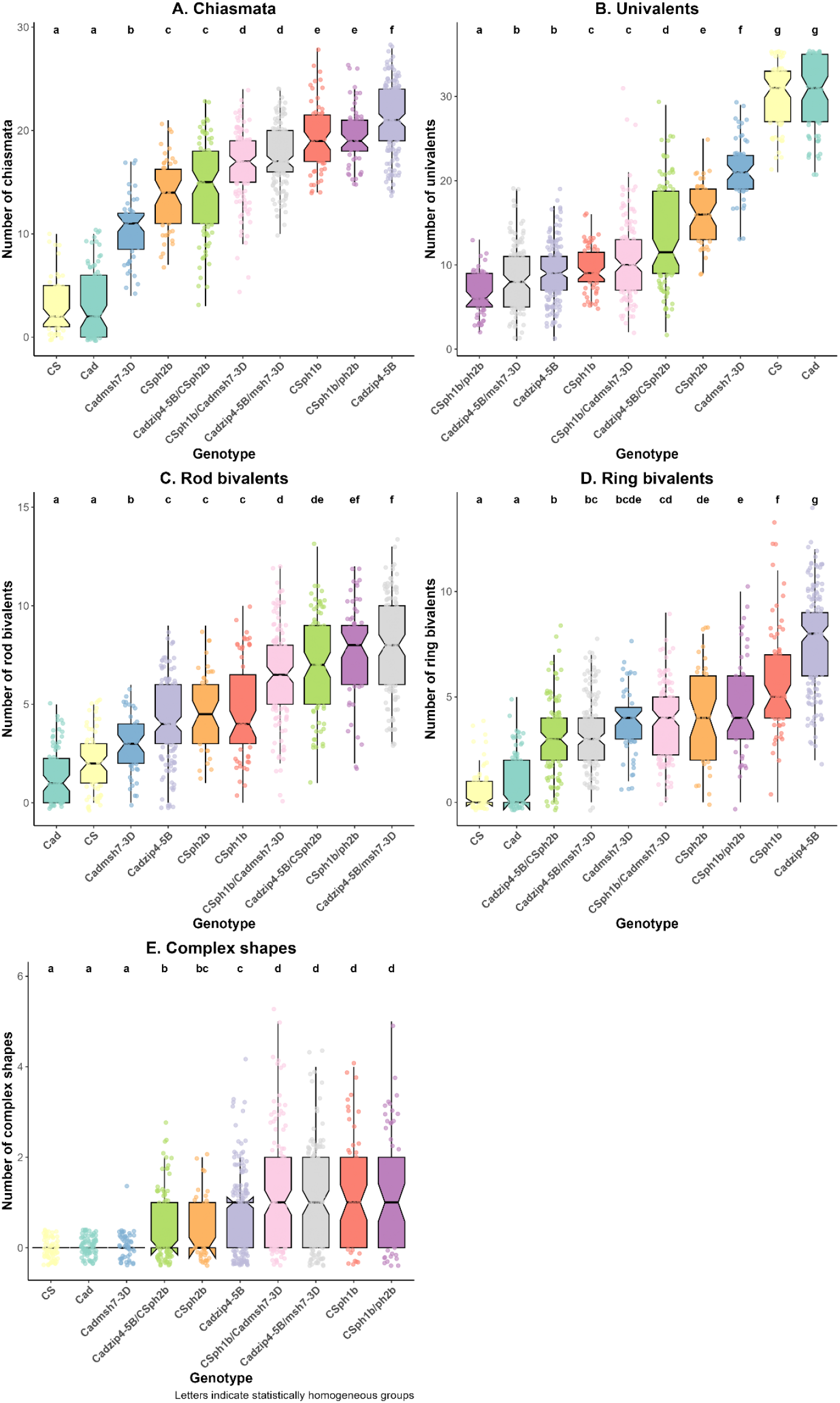
Boxplots showing the numbers of chiasmata (A), univalents (B), bivalents (C: rod and D: ring) and multivalents (E), according to genotypes at the *Ph* loci in Cadenza (Cad) or Chinese Spring (CS) backgrounds in the interspecific hybrids. Boxplots illustrate the distribution of each meiotic configuration at metaphase I. Pairwise comparisons between genotypes were assessed using the Mann–Whitney test. Based on the resulting p-values, genotypes were grouped according to statistical significance. Letters above the boxplots indicate significance groups: genotypes sharing the same letter are not significantly different, while those with different letters show statistically significant differences.

Hybrids derived from crosses involving wild-type plants gave similar values for CS and Cad, with mostly univalents (around 30; Fig. 2B) and only occasional bivalents (Fig. 2C-D), resulting in few chiasmata (3.25 ± 3.22 for Cad and 3 ± 2.46 for CS; Fig. 2A). This indicates that control of homoeologous CO was comparable between the two cultivars.

In contrast, when comparing the *ph2b* mutant in CS and the *msh7-3D* mutation in Cad, clear differences in meiotic configuration were observed. Univalent number was significantly higher in Cad *msh7-3D* (21.78 ± 3.55) compared to CS *ph2b* (15.94 ± 3.47; Fig. 2B). Bivalent number was higher in CS *ph2b* (4.73 ± 1.85 rods and 4.08 ± 2.11 rings) than in Cad *msh7-3D* (2.88 ± 1.41 rods and 3.71 ± 1.6 rings; Fig.2C and D). Cad *msh7-3D* and CS *ph2b* both showed very few multivalents (0.02 ± 0.14 and 0.48 ± 0.65 respectively; Fig. 2E). As a result, the total number of chiasmata was significantly higher in CS *ph2b* compared to Cad *msh7-3D* (13.88 ± 3.47 and 10.31 ± 3.07 respectively; Fig. 2A).

Different conclusions were drawn from the single *ph1* mutations (CS *ph1b* or Cad *zip4-5B*). As expected, both Cad *zip4-5B* and CS *ph1b* genotypes displayed fewer univalents and more bivalents, resulting in more chiasmata compared to controls and to the *ph2b* or *msh7-3D* single mutants. Interestingly, the numbers of rod bivalents were not significantly different between Cad *zip4-5B* and CS *ph1b* (4.16 ± 1.93 and 4.78 ± 2.35 respectively; Fig. 2C). In contrast, significantly more ring bivalents were found with Cad *zip4-5B* compared to CS *ph1b* (7.7 ± 2.3 versus 5.85 ± 2.52; Fig. 2D), whereas for multivalents, the converse was true (0.73 ± 0.87 versus 1.25 ± 1.16; Fig. 2E). This led to a significant reduction in univalent number, especially in Cad *zip4-5B* (8.85 ± 3.47 in Cad *zip4-5B* and 9.69 ± 2.6 in CS *ph1b*; Fig. 2B), as well as an increase in total chiasmata (21.21 ± 3.33 in Cad *zip4-5B* and 19.4 ± 3.17 in CS *ph1b*; Fig. 2A).

### 3.2 Analysis of interspecific hybrids combining *ph1* and *ph2* mutations at meiotic metaphase I

To further our analysis, both mutations were combined, either in the same wheat genetic background or in a recombinant genome. When both *ZIP4-5B* and *MSH7-3D* were mutated in the same genetic background, Cad *zip4-5B/msh7-3D* showed significantly more univalents than CS *phb1/ph2b* (8.4 ± 3.8 and 6.63 ± 2.53; Fig. 2B) and significantly fewer ring bivalents (3.37 ± 1.72 and 4.56 ± 2.16; Fig. 2C), resulting in a lower number of chiasmata (17.54 ± 2.92 and 19.63 ± 2.89; Fig. 2A).

Recombinant double mutants, also showed significant differences. Univalents were more frequent in Cad *zip4-5B*/CS *ph2b* (13.29□±□5.95) compared to CS *ph1b*/Cad *msh7-3D* (10.79□±□4.82). The two genotypes had comparable numbers of rod bivalents, but CS *ph1b*/Cad *msh7-3D* presented a higher frequency of ring bivalents (3.75 ± 1.71 vs 3.11 ± 1.87) and multivalents (1.13□±□1.23 vs. 0.46□±□0.72). Consequently, Cad *zip4-5B*/CS *ph2b* exhibited fewer chiasmata (14.32□±□4.4) than CS *ph1b*/Cad *msh7-3D* (16.69□±□3.29).

Interestingly, the recombinant mutants Cad *zip4-5B*/CS *ph2b* and CS *ph1b*/Cad *msh-3D* exhibited fewer chiasmata and more univalents than the non-recombinant genotypes (Cad *zip4-5B/msh7-3D* and CS *ph1b/ph2b*), suggesting the involvement of other interacting factors at the whole-genome level affecting the efficiency of these mutations on homoeologous CO. All double mutants had a reduced CO level compared to the *ph1* single mutants, apart from the CS *ph1b/ph2b* double mutant which had a similar level of COs to the single CS *ph1b* mutant. Thus, there was no additive effect of the two mutations on homoeologous CO; in fact, there was a tendency towards a reduction in CO frequency. This aligns with the reduction in ring bivalents and increase in rod bivalents observed with the double mutants.

### 3.3 Meiotic dynamics in wheat and interspecific hybrids *ph1* and *ph2* single or double mutants

Whilst collecting material to study the progression of meiosis in the interspecific hybrids, we observed considerable variation in meiotic behaviour between individuals. In some genotypes, anther size differed within the same flower (individuals 05B, 10D, 20F, 23F, 25G, 27G, 14E in Supplementary Table 1; Supplementary Figure 3), or were even atrophied and white (individuals 10D, 20F, 25G). Anthers were sometimes desynchronized (individuals 05B, 10D, 20F, 23F, 25G, 27G, 14E) even within the same floret, with meiotic sacs at different stages (individuals 19E, 26G, 28G) or with meiocytes of different size. To investigate this further, we conducted a more detailed analysis of meiosis in some of these interspecific hybrids in order to visualize all types of anomalies encountered. We selected five interspecific hybrids carrying single or double mutations: mutant 04B (CS *ph2b*); 07C (Cad *msh7-3D*); 23F (CS *ph1b*/Cad *msh7-3D*); 28G (Cad *zip4-5B*/CS *ph2b*); 17E (Cad *zip4-5B/msh7-3D*) in Supplementary Table 1. A CS wildtype bread wheat (WT CS) was included as control. We then followed their progression of meiosis, which clearly revealed meiotic disturbances in the interspecific hybrids derived from mutants (Figure 3).

**Figure 3.**
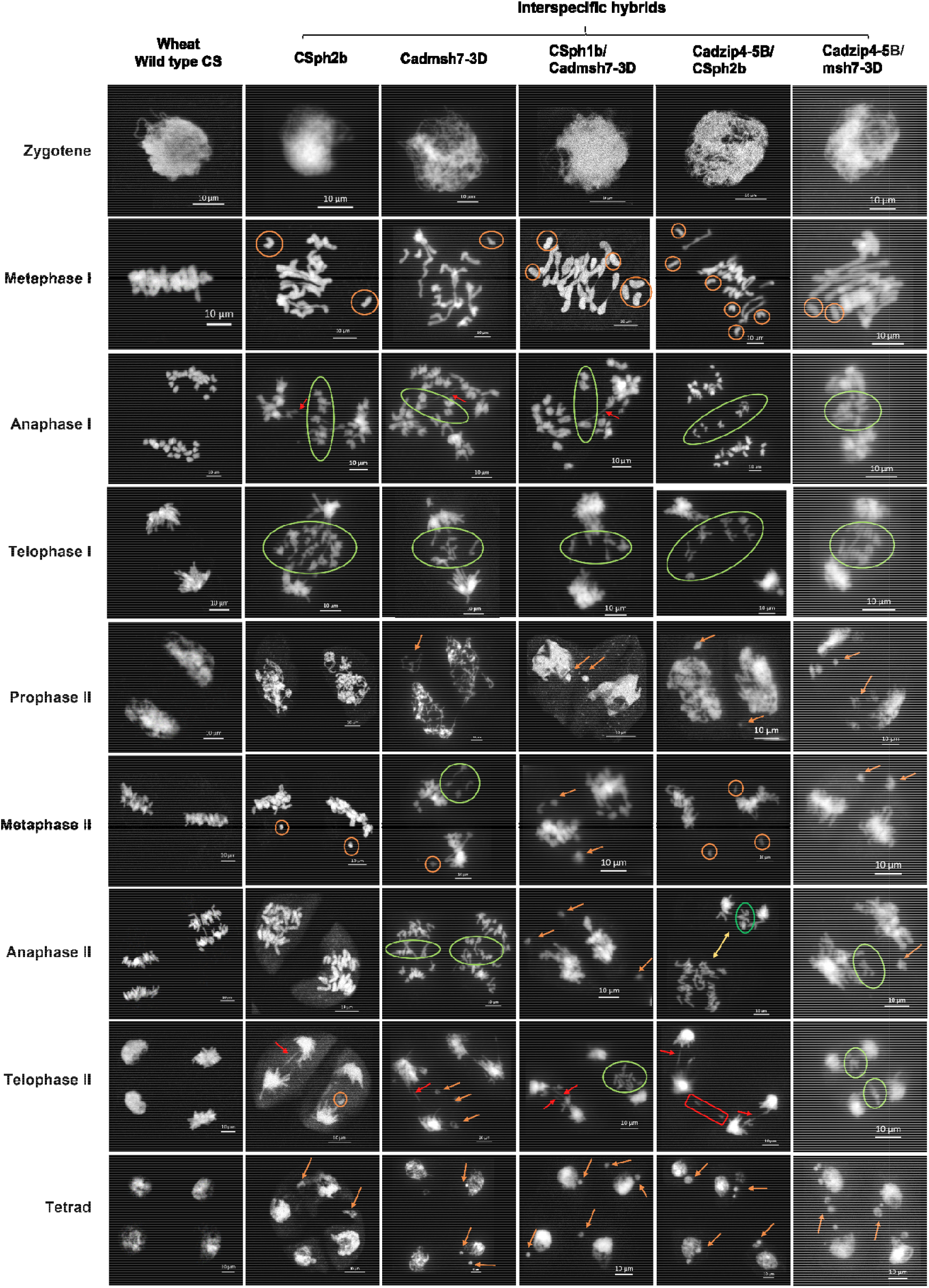
Meiotic progression in control wheat (WT CS) and selected interspecific hybrids after DAPI staining. Numerous disturbances during meiosis were observed in all interspecific hybrids. Symbols indicate specific abnormalities: orange circle, univalents; green circle, segregation anomalies; orange arrow, chromosome fragmentation; red arrow, bridges; yellow arrow, desynchronization of meiotic stages between the two cellular compartments; red rectangle, structural disorganization of a cellular compartment. Meiocytes were observed under a microscope with a 40× objective; scale bar, 10 µm.

In the interspecific hybrid Cad *zip4-5B*/CS *ph2b*, meiocytes of different sizes were observed throughout all meiotic phases, whereas in the Cad *zip4-5B/msh7-3D*, small meiocytes were only observed from prophase II. From prophase I, fragmentations were detected during the diplotene and diakinesis stages in the interspecific hybrids Cad *msh7-3D*, CS *ph1b*/Cad *msh7-3D*, and Cad *zip4-5B/msh7-3D* (Supplementary Figure 4). Additional fragmentations were observed from prophase II in most interspecific hybrids, except for CS *ph2b*, which only showed fragmentations during the tetrad stage. In metaphase I, one or more univalents were visible, and in anaphase I, anaphase II and telophase II, bridges were observed in all individuals, as well as segregation irregularities during anaphase I, telophase I, metaphase II and telophase II. Moreover, desynchronization of the meiotic stage and structural disorganization between the two cell compartments were specifically observable in anaphase II and telophase II in Cad *zip4-5B*/CS *ph2b*.

Thus, the Cad *zip4-5B*/CS *ph2b* interspecific hybrid appears to be the most affected among all. It not only displays anthers of variable sizes and a desynchronization of the meiotic stage (anaphase II; Figure 3), but also exhibits chromosomal exchanges between two cellular compartments. This phenomenon was observed exclusively in this hybrid, indicating a disruption in the structural organization of the cellular compartments during telophase II (Figure 3). Moreover, among all hybrids, the Cad *zip4-5B*/CS *ph2b* shows the highest percentage of anomalies observed at the tetrad stage (37%), including 23% triads, 13% polyads, and 1% dyads (Supplementary Table 5).

## 4 Discussion

For nearly 40 years, wheat breeders have used the Sears mutant, carrying a deletion of the *Ph1* locus, to introgress agronomically useful genes or alleles (homoeologous chromosomes) from wild relatives into wheat (King et al., 2019). Recently, researchers revealed that the effect of the *Ph1* locus in tetraploid and hexaploid wheat (consisting of both homologous and homoeologous chromosomes) is explained by a dual function *ZIP4-5B* gene, coding for a protein required in type-I CO formation (Chelysheva et al., 2007). This promotes the synapsis of homologous chromosomes during meiosis, whilst also suppressing crossover between the homoeologous chromosomes (Draeger et al., 2023; Martín et al., 2021, 2017; Rey et al., 2017).

As regards to *Ph2*, this locus has also been recognised as playing a role in CO during wheat meiosis, its function has only recently been attributed to the *MSH7-3D* gene (Serra et al., 2021), coding for the plant specific MSH7 protein (*Mut-S homolog 7*). This protein belongs to the DNA Mismatch Repair (MMR) complex (Culligan and Hays, 2000). It has been suggested that this MSH7-3D protein could play a crucial role during the early steps of recombination, by rejecting heteroduplex single-strand invasion between homoeologues (Serra et al., 2021).

Our work investigated whether the wheat genetic background and the combinations of different mutations in these two genes could enhance the efficiency of crossover between homoeologous chromosomes in hybrids of wheat and a wild relative. In our study, hybrids derived from crosses between *Ae. variabilis* (Aev) and Chinese Spring (CS) *ph1b* and/or *ph2b* mutants developed in the 1970s, were compared to hybrids derived from crosses using the recent Cadenza (Cad) *zip4-5B* and/or *msh7-3D*.

### 4.1 Wheat genetic background and/or mutation type in *Ph1* and *Ph2* affects the level of homoeologous recombination

In interspecific hybrids derived from crosses between different bread wheat (CS and Cad) and the wild tetraploid species Aev, we demonstrated that CS and Cad wild types gave similar results, both displaying similarly high numbers of univalents and very low chiasma frequencies, indicating a comparable suppression of homoeologous CO in both wheat backgrounds. Additionally, we showed that *ph1* and *ph2* mutations result in a global increase of rod and ring bivalents and multivalents at the expense of univalents, leading to an increased number of chiasmata between homoeologous chromosomes. We also observed that the effects of *ph1* mutations alone are greater than those of *ph2*, regardless of whether the background is Cad or CS. For example, in terms of chiasma number, the effect of the *ph1* mutation was approximately two-fold higher than the *msh7-3D* mutation in a Cad background, and about 1.4 times higher than the *ph2b* mutation in a CS background.

All these findings are in line with previous studies (Table 2; Bochev et al., 1978; Ceoloni and Donini, 1993; Rey et al., 2017; Serra et al., 2021).

**Table 2:** Comparative analysis of number of chiasmata observed in this study with values extracted from literature.

| Type of cross | Shapes | Our study |  | Other references |  |
| --- | --- | --- | --- | --- | --- |
|  |  | Chinese Spring | Cadenza | Chinese Spring | Cadenza |
| **WT X *Ae. variabilis*** | *Multivalents* | 0.00 | 0.00 | 0.00 a | 0.00 a |
|  | *Bivalents* | 2.50 ± 1.82 | 2.39 ± 2.11 | 1.38 ± 0.12 b | 1.1 ± 0.09 c |
|  | *Univalents* | 30.00 ± 3.65 | 30.23 ± 4.22 | 32.15 ± 0.25 b | 32.05 ± 0.24 d |
| ***ph1* X *Ae. variabilis*** | *Multivalents* | 1.25 ± 1.16 | 0.73 ± 0.87 | 2.74 ± 0.11 e | 1.3 ± 0.14 d |
|  | *Bivalents* | 10.63 ± 2.33 | 11.86 ± 2.20 | 7.56 ± 0.11 e | 8.01 ± 0.25 d |
|  | *Univalents* | 9.69 ± 2.60 | 8.85 ± 3.47 | 11.06 ± 0.18 e | 14.74 ± 0.29 d |
| ***ph2* X *Ae. variabilis*** | *Multivalents* | 0.48 ± 0.65 | 0.02 ± 0.14 | 0.04 ± 0.04 b | 0.12 ± 0.03 c |
|  | *Bivalents* | 8.81 ± 1.78 | 6.59 ± 1.75 | 3.34 ± 0.24 b | 5.55 ± 0.19 c |
|  | *Univalents* | 15.94 ± 3.47 | 21.78 ± 3.55 | 27.26 ± 0.48 b | 22.86 ± 0.09 c |

a: all references; b: Bochev et al. 1978; c: Serra et al., 2021; d: Rey et al. 2017; e: Ceoloni and Donini 1993.

Interestingly, differences were observed between Cad *zip4-5B* and CS *ph1b* mutants. Although both genotypes displayed a similar number of rod bivalents, Cad *zip4-5B* formed a higher number of ring bivalents, whereas CS *ph1b* showed a greater frequency of multivalents. This suggests that Cad *zip4-5B* supports a more regular pairing behaviour. This could be explained by the fact that CS *ph1b* has accumulated rearrangements during its propagation, which will lead to the background occurrence of multivalents (Martín et al., 2018). However, the difference between the two mutants is not large, suggesting that the EMS mutation of *zip4-5B* in Cadenza mimics well the effect of the deletion of *Ph1* in Chinese Spring.

We also observed consistent values for the Cad *msh7-3D* mutant compared to previous studies. However, in our study, the CS *ph2b* mutant exhibited more bivalents and multivalents, and therefore more chiasmata, compared to the Cad *msh7-3D* mutant. It is possible, as in the case of the CS *ph1b* mutant, that the forty-year-old CS *ph2b* mutant also carries background chromosome rearrangements, affecting its meiotic configurations in hybrids. Furthermore, our study highlights that the Chinese Spring double mutant has a higher level of CO than the Cadenza double mutant. This may again reflect the relative level of background rearrangements in the older CS *ph1b* and CS *ph2b* compared to the very recent Cad *zip4-5B* and Cad *msh7-3D* mutants.

### 4.2 Accumulation of *ph1* and *ph2* mutations does not improve homoeologous recombination

We observed no further increase in chiasmata number in the double mutants compared to the *ph1b* or *zip4-5B* single mutants alone; in some cases, chiasma number was slightly lower. This indicates that when both *zip4-5B* and *msh7-3D* are mutated/eliminated, homoeologous CO reaches its maximum with the *ph1* mutation alone, as Mello-Sampayo and Canas (1973) reported in the context of CS x *Ae. sharonensis*. This evidence suggests that the two genes may be acting within the same pathway (type I CO), albeit at different stages of the process (Sourdille et al., 2025). Right after double-strand breaks are produced in early prophase I, it is likely that their repair can be initiated on both homologues or homoeologues that are very close because of the bouquet structure (Martinez et al., 2001; Osman et al., 2011). However, the mismatch repair protein complex TaMSH2/TaMSH7 (*Ph2*) certainly detects the numerous differences between the homoeologues and contributes to prevent the single-strand invasion and the initiation of homoeologous synapsis (Serra et al., 2021). In the same time, *TaZip4-B2* (*Ph1*) promotes homologous synapsis during the telomere bouquet stage (Martín et al., 2014). However, it is likely that not all homoeologous invasions are removed, especially in regions of extremely high similarity between homoeologues (*i*.*e*. at the gene level). In these rare cases, the *Ph1* locus acts also downstream by preventing the maturation of crossovers between homoeologous chromosomes (Martín et al., 2014). This ultimately contributes to the establishment of a diploid-like meiotic behaviour of bread wheat.

Thus, homoeologous COs are thought to arise predominantly via the class I pathway (ZMM dependent), which exhibits interference and is more tolerant to sequence polymorphisms than the class II pathway (MUS81 dependent; Ziolkowski, 2023). Indeed, class II crossovers preferentially occur in homozygous regions (Blackwell et al., 2020; Crismani et al., 2012; Fernandes et al., 2017), whereas class I crossovers can form in polymorphic regions and are redistributed according to heterozygosity patterns (Blackwell et al., 2020; Ziolkowski et al., 2015). Consistently, genes promoting class I COs have been directly implicated in the regulation of homoeologous recombination as in *Brassica napus* (Gonzalo et al., 2019; Grandont et al., 2014), supporting the idea that most homoeologous COs arise through the class I pathway rather than the class II pathway.

Although double mutants do not increase the number of chiasmata, they all exhibit a greater number of rod bivalents and fewer ring bivalents than the *ph1b* or *zip4-5B* single mutants, suggesting a shift in the type of chromosome associations during meiosis. Loss of ring bivalents for rod bivalents is often associated with problems during synapsis. It has been shown that homoeologous synapsis can only occur after the telomere bouquet stage has completed, and not during the bouquet stage (Martín et al., 2017). Homologous synapsis, in contrast, can occur during the bouquet stage (Martín et al., 2017). The combined loss of *ZIP4-5B* and *MSH7-3D* may affect the synapsed homoeologues in some way at this late stage, resulting in early resolution of homoeologues as rod bivalents rather than ring bivalents. It has already been shown that *ZIP4-5B* is required for completion of synapsis (Draeger et al., 2023).

Even though we did not observe any additive effect between *zip4-5B* (*ph1*) and *msh7-3D* (*ph2*) mutants in our study, additive interactions have been reported in other *Ph* combinations in crosses involving Chinese Spring x *Aegilops variabilis*. Liu et al., (2003) demonstrated that association of the *phKL* locus with *ph2* leads to a significant 40% increase in chiasma number, primarily due to an increase in rod bivalents. However, this combination is 50% less effective in terms of chiasma number compared to Chinese Spring *ph1b* mutants x *Aegilops variabilis*. Thus, it is the single mutation of *ZIP4-5B* that appears to maximize the number of chiasmata in this context. However, combinations with other mutations, possibly focusing on proteins involved in class II COs, could contribute to optimizing the level of homoeologous crossover, resulting in more efficient introgression from wild relatives into wheat.

Interestingly, the combination of *zip4-5B* (*ph1*) and *msh7-3D* (*ph2*) mutations was more efficient when they were in the same genetic background (either Cad or CS), compared to when they were present in a Cad/CS recombinant background. The number of univalents was twice as high in Cad *zip4-5B*/CS *ph2b* than in CS *ph1b*/*ph2b* (13.29 ± 5.95 and 6.63 ± 2.53 respectively) and 1.6 times higher in CS *ph1b*/Cad *msh7-3D* (10.79 ± 4.82). The same was true albeit lower with Cad *zip4-5B/msh7-3D* (8.4 ± 3.8), which showed 1.6 and 1.3 fewer univalents respectively, compared to Cad *zip4-5B*/CS *ph2b* and CS *ph1b*/Cad *msh7-3D*. Moreover, recombinant double mutant hybrids (notably Cad *zip4-5B*/CS *ph2b*) exhibited increased meiotic disruptions. This suggests that there are other factors elsewhere in the genome interacting with *ZIP4-5B* and/or *MSH7-3D* to control homoeologous CO, and that these interactors are more efficient when coming from the same background. For example, the MSH7 protein is working as a heterodimer with MSH2 to form a complex that drives mismatch repair (Culligan and Hays, 2000; Wu et al., 2003). It is likely that MSH2 and MSH7 could have evolved concomitantly in different backgrounds to retain maximal efficiency. Different alleles exist for *MSH7-3A* and *MSH7-3B* (Serra et al., 2021), and this could be the same for MSH2, as well as for other factors that could be interacting with *ZIP4-5B* and/or *MSH7-3D*.

*Ph1* has always been assumed to be the main regulator of homoeologous CO in wheat. However, the homoeologous copies of *Msh7-3D* (*Ph2*) on chromosomes 3A and 3B may correspond to other regulators of homoeologous CO identified previously (Driscoll, 1972; Mello-Sampayo and Canas, 1973; Miller et al., 1983). It is likely that if mutants for two or three copies of MSH7 were to be produced, the effect on homoeologous CO could be greater. This is in accordance with previous observations showing that in mutants losing both 3AS (*Ph3*) and 3DS (*Ph2*), chiasma frequency between homoeologous chromosomes is similar to that caused by the deficiency of *Ph1* alone (Driscoll, 1972). Also, other genes such as *Asy1* (Di Dio et al., 2023) or *RECQ4* (Bazile et al., 2024) seem to affect homoeologous CO. It is likely that additional genes will be discovered in the near future.

### 4.3 Meiosis is disrupted in *ph1* and *ph2* mutants

Progression of meiosis was clearly disrupted in all our interspecific hybrids. Similar disruptions have been reported in previous studies, notably in interspecific hybrids between *Triticum aestivum* and *Agropyron cristatum*, which revealed a series of major meiotic disruptions (Limin and Fowler, 1990). In our material, we observed chromosome fragmentation and bridges, most likely resulting from homoeologous COs that were not properly resolved during meiosis, due to the absence of the *ZIP4-5B* and *MSH7-3D* proteins.

These abnormalities also stem from complex interactions between genomes from distinct parental species (i.e. wheat and *Ae. variabilis*), which can lead to segregation distortion (Endo, 1990). Such genomic interactions can generate various disruptive elements, such as genetic imbalances, chromosomal incompatibilities, abnormalities in chiasma formation, divergent hormonal responses, epigenetic alterations and variations in chromosome number. Rooted in genetic, structural and regulatory differences between parental species, these factors are likely to compromise orderly chromosome segregation and the process of genetic recombination. This highlights the importance of controlling homologous and homoeologous recombination to prevent loss of fertility in the progeny.

## 5 Conclusion

The production of F1 interspecific hybrids between wheat mutants (*ph1* and/or *ph2*) and *Aegilops variabilis* provides a valuable pre-breeding resource, as it enhances homoeologous recombination and facilitates the introgression of useful traits from *Aegilops* species that are otherwise difficult to access through conventional crosses due to meiotic barriers. Following the restoration of fertility in the F1 interspecific hybrid, successive backcrosses with elite wheat are required to stabilize the genome and facilitate the evaluation of successfully introgressed *Aegilops* segments.

Here, we demonstrate that accumulation of *ph1* (*zip4-5B*) and *ph2* (*msh7-3D*) mutations do not enhance homoeologous recombination in comparison with the *ph1* alone. Thus, we can confirm that the *ph1* mutation remains the most efficient way to perform such introgressions within the wheat genome. However, its strong effect is accompanied by considerable genomic instability, including multivalent formation and abnormal chromosome segregation, especially in the deleted mutants (Alabdullah et al., 2021; Sánchez-Morán et al., 2001; Sourdille et al., 2025). This instability often results in increased aneuploidy and a loss of fertility, raising concerns about the persistence of deleterious rearrangements in advanced lines. In contrast, loss of *Ph2* function induces a moderate yet significant increase in homoeologous crossovers while it could maintain better overall genome integrity and fertility, making it a more practical option for breeders seeking to minimize post-introgression disruptions (Serra et al., 2021).

The choice of genetic background is also a critical factor in wheat pre-breeding programs. Chinese Spring is widely employed as the reference background for mutations, but most of the available mutants correspond to old deletion lines, and the CS background itself is limited by poor agronomic performance. The use of the CS background will necessitate that several backcrosses are carried out to achieve agronomic performance levels comparable to modern elite germplasm (Türkösi et al., 2022). In contrast, the Cadenza background is preferred for breeding because it exhibits good agronomic performance (Ma et al., 2015) and, with EMS-derived point mutants, could result in lower meiotic instability compared to older CS deletion lines.

Ultimately, transferring or generating *Ph* mutations directly in elite wheat cultivars could further streamline breeding pipelines by reducing the need for extensive backcrossing and accelerating the release of improved varieties (Li et al., 2020; Türkösi et al., 2022).

## 7 Author Contributions

PS, GM, CJB, designed the experiments and raised funding; CH, IN, ACM, M-DR, ABB, realised all experiments; CH, IN, ABB, analysed meiotic phenotypes; CH did all statistics, figures and tables; CH and PS drafted the manuscript; all authors corrected and approved the final version of the manuscript.

## 9 Acknowledgments

The Authors thank Isabelle LHOMMET for her help during sample collection and Ludovic GEORGES for genotyping the interspecific hybrids. Members of the VégéPôle platform are also acknowledged for their help during plant growth period as well as the members of GENTYANE platform for their support for molecular analyses.

