## Supplementary material for "Accumulation of *ph1* (*zip4-5B*) and *ph2* (*msh7-3D*) mutations fails to boost homoeologous recombination in hexaploid wheat"

1. **Supplementary Table 1:** Classification of the 29 hybrids according to their origin, *Ph1* and *Ph2* loci genotyping and selection for phenotyping.

| ***Group*** | ***Name*** | ***Origins*** | ***Genotyping*** | ***Phenotyping*** |
| --- | --- | --- | --- | --- |
| Control | 01A | (CS x Cad) x Aev | CS Cad |  |
|  | 02A | (CS x Cad) x Aev | CS Cad |  |
|  | 03A | (CS x Cad) x Aev | CS Cad |  |
| Single mutant | 04B | (CS ph2b x Cad 2006) x Aev | CS ph2b | x |
|  | 05B | (CS ph2b x Cad 2006) x Aev | CS ph2b |  |
|  | 06B | (CS ph2b x Cad 2006) x Aev | CS ph2b |  |
|  | 07C | (CS ph2b x Cad 2006) x Aev | Cad msh7-3D |  |
|  | 08C | (CS ph2b x Cad 2006) x Aev | Cad msh7-3D | x |
|  | 09C | (CS ph2b x Cad 2006) x Aev | Cad msh7-3D |  |
| Double mutated parental CS | 10D | (CS ph1b ph2b x Cad 0348 2006) x Aev | CS ph1b/CS ph2b | x |
|  | 11D | (CS ph1b ph2b x Cad 0348 2006) x Aev | CS ph1b/CS ph2b |  |
|  | 12D | (CS ph1b ph2b x Cad 0348 2006) x Aev | CS ph1b/CS ph2b |  |
|  | 13D | (Cad 1691 2006 x CS ph1b/ph2b) x Aev | CS ph1b/CS ph2b | x |
| Double mutated parental Cad | 14E | (CS ph1b ph2b x Cad 0348 2006) x Aev | Cad zip4-5B/Cad msh7-3D |  |
|  | 15E | (CS ph1b ph2b x Cad 0348 2006) x Aev | Cad zip4-5B/Cad msh7-3D | x |
|  | 16E | (CS ph1b ph2b x Cad 0348 2006) x Aev | Cad zip4-5B/Cad msh7-3D |  |
|  | 17E | (CS ph1b ph2b x Cad 0348 2006) x Aev | Cad zip4-5B/Cad msh7-3D | x |
|  | 18E | (CS ph1b ph2b x Cad 0348 2006) x Aev | Cad zip4-5B/Cad msh7-3D |  |
|  | 19E | (CS ph1b ph2b x Cad 0348 2006) x Aev | Cad zip4-5B/Cad msh7-3D | x |
| Double mutant recombinant CS-Cad | 20F | (CS ph1b ph2b x Cad 0348 2006) x Aev | CS ph1b/Cad msh7-3D |  |
|  | 21F | (CS ph1b ph2b x Cad 0348 2006) x Aev | CS ph1b/Cad msh7-3D | x |
|  | 22F | (CS ph1b ph2b x Cad 0348 2006) x Aev | CS ph1b/Cad msh7-3D | x |
|  | 23F | (Cad 1691 2006 x CS ph1b ph2b) x Aev | CS ph1b/Cad msh7-3D | x |
|  | 24F | (Cad 1691 2006 x CS ph1b ph2b) x Aev | CS ph1b/Cad msh7-3D |  |
|  | 25G | (CS ph1b ph2b x Cad 0348 2006) x Aev | Cad zip4-5B/CS ph2b |  |
|  | 26G | (CS ph1b ph2b x Cad 0348 2006) x Aev | Cad zip4-5B/CS ph2b |  |
|  | 27G | (CS ph1b ph2b x Cad 0348 2006) x Aev | Cad zip4-5B/CS ph2b | x |
|  | 28G | (CS ph1b ph2b x Cad 0348 2006) x Aev | Cad zip4-5B/CS ph2b |  |
|  | 29G | (CS ph1b ph2b x Cad 0348 2006) x Aev | Cad zip4-5B/CS ph2b | x |

2. **Supplementary Table 2:** Classification of the 33 hybrids used in the study

| ***Group*** | ***Made in*** | ***Origins*** | ***Genotyping at the ph loci*** | ***Number of plants*** | ***Name of plants at INRAE*** |
| --- | --- | --- | --- | --- | --- |
| Control | JIC | Cad x Aev | Cad | 5 |  |
|  | JIC | CS x Aev | CS | 2 |  |
| Single mutant | JIC | Cad 0348 x Aev | Cad zip4-5B | 5 |  |
|  | JIC | Cad 1691 x Aev | Cad zip4-5B | 7 |  |
|  | JIC | CS ph1b x Aev | CSph1b | 2 |  |
|  | INRAE | (CS ph2b x Cad 2006) x Aev | Cad msh7-3D | 1 | 08C |
|  | INRAE | (CS ph2b x Cad 2006) x Aev | CSph2b | 1 | 04B |
| Double mutant CS | INRAE | (CS ph1b ph2b x Cad 0348 2006) x Aev | CS ph1b/CS ph2b | 1 | 10D |
|  | INRAE | (Cad 1691 2006 x CS ph1b ph2b) x Aev | CS ph1b/CS ph2b | 1 | 13D |
| Double mutant Cad | INRAE | (CS ph1b ph2b x Cad 0348 2006) x Aev | Cad zip4-5B/Cad msh7-3D | 3 | 15E, 17E, 19E |
| Double mutant recombined Cad/CS | INRAE | (CS ph1b ph2b x Cad 0348 2006) x Aev | CS ph1b/Cad msh7-3D | 2 | 21F, 22F |
|  | INRAE | (Cad 1691 2006 x CS ph1b ph2b) x Aev | CS ph1b/Cad msh7-3D | 1 | 23F |
|  | INRAE | (CS ph1b ph2b x Cad 0348 2006) x Aev | Cad zip4-5B/CS ph2b | 2 | 27G, 29G |

3. **Supplementary Table 3:** Primer sequences used for *Ph* gene tracking in CS and Cad background.

| **Name of primers** | **Sequence 5'-3'** | **Tm** | **Amplification product** |
| --- | --- | --- | --- |
| Cad_0348_F-wildtype | TTAGCCACCCTTCTACCCATATTC | 60°C | 50 bp |
| Cad_0348_F-mutant | TTAGCCACCCTTCTACCCATATTT |  |  |
| Cad_0348_R-common | CTGAACTGGTTTGCAGGCAATA |  |  |
| Cad_1691_F-wildtype | GGGCGATCAGTCCCTCGC | 60°C | 86 bp |
| Cad_1691_F-mutant | GGGCGATCAGTCCCTCGT |  |  |
| Cad_1691_R-common | GATCTGGTATACTTGCGGGG |  |  |
| Cad_2006_F-common | CTATTCTCATCTGGTTTCTTTGT | 55°C | 81 bp |
| Cad_2006_R-wildtype | CCAGGTAGTGGGGTCAGTAG |  |  |
| Cad_2006_R-mutant | CCAGGTAGTGGGGTCAGTAA |  |  |
| CS_ph1b_F | TATGGGTAGAAGGGTGGCT | 60°C | 250 bp |
| CS_ph1b_R | GTTTACCTTGCCAGCTCTG |  |  |
| CS_ph2b_F | AAGCTAGAGGACCAAATTCAA | 55°C | 1240 bp |
| CS_ph2b_R | TAGAGTAACCTGAAAATTCCAACAAC |  |  |

To follow Cad0348 or Cad1691:

- 1 well: primer Cad_0348_F-wildtype + primer Cad_0348_R-common OR primer Cad_1691_F-wildtype + primer Cad_1691_R-common
- 1 other well: primer Cad_0348_F-mutant + primer Cad_0348_R-common OR primer Cad_1691_F-mutant + primer Cad_1691_R-common

If an individual shows amplification for the wild-type primers only, they are homozygous **Cad *Ph1/Ph1***. If amplification occurs only for the mutant primers, the individual is homozygous mutant **Cad *ph1/ph1***. If amplification is detected for both primers, the individual is heterozygous **Cad *Ph1/ph1***.

This is what the markers give on the parents of our experiment:

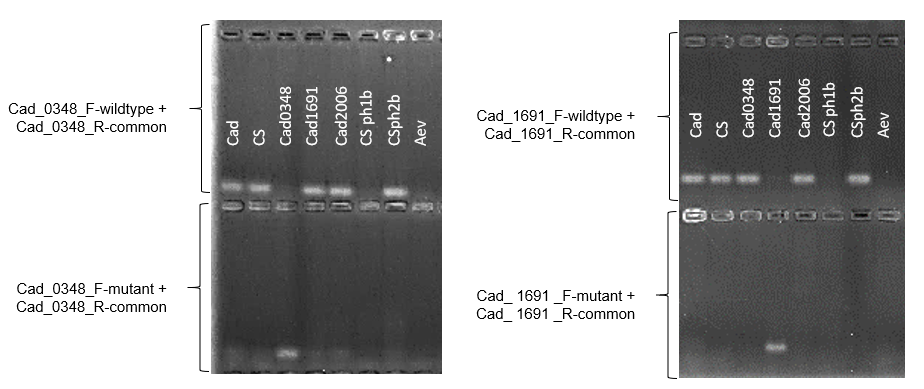

To follow Cad2006:

- 1 well: primer Cad_2006_R-wildtype + primer Cad_2006_F-common
- 1 other well: primer Cad_2006_R-mutant + primer Cad_2006_F-common

If an individual shows amplification for the wild-type primers only, they are homozygous **Cad *Ph2/Ph2***. If amplification occurs only for the mutant primers, the individual is homozygous mutant **Cad *ph2/ph2***. If amplification is detected for both primers, the individual is heterozygous **Cad *Ph2/ph2***.

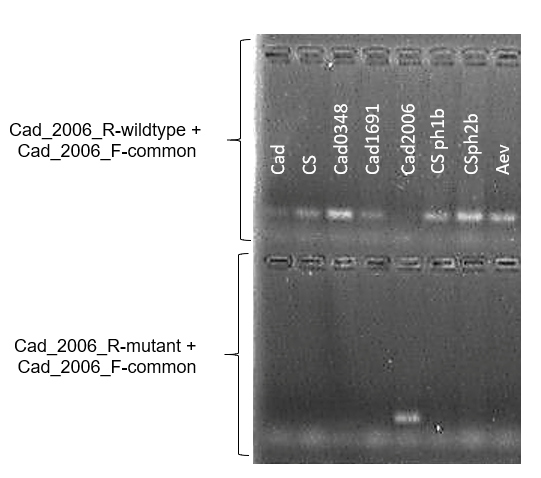

To follow CSph1b:

- 1 well: primer CS_ph1b_F + CS_ph1b_R

If an individual shows amplification, they are homozygous wild type **CS *Ph1/Ph1*** or heterozygous **CS *Ph1/ph1***. If no amplification is observed, the individual is homozygous mutant **CS** ***ph1/ph1***.

To follow CSph2b:

- 1 well: primer CS_ph2b_F + CS_ph2b_R

If an individual shows amplification, they are homozygous mutant **CS *ph2/ph2*** or heterozygous **CS *Ph2/ph2***. If no amplification is observed, the individual is homozygous wild type **CS** ***Ph2/Ph2***.

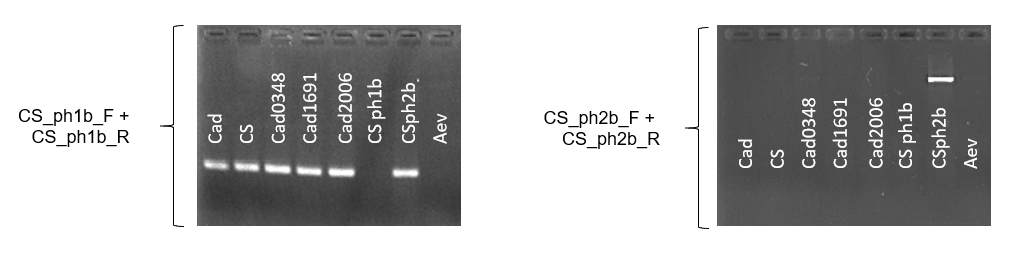

4. **Supplementary Figure 1:** Types of chromosomal pairing structures and corresponding chiasmata numbers.

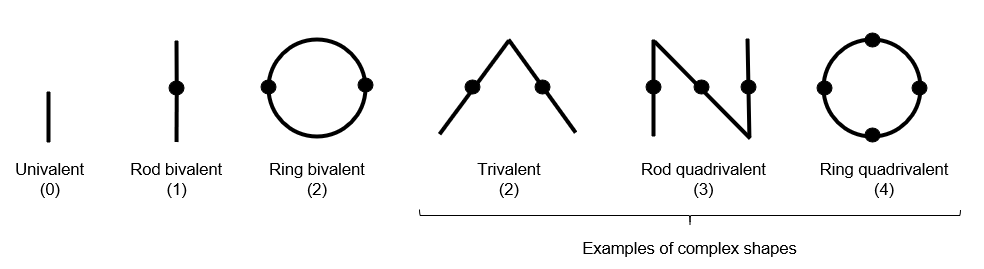

Various chromosomal pairing configurations can be observed during meiosis, including univalent, rod bivalent, ring bivalent, and multivalent (trivalent, rod quadrivalent, and ring quadrivalent formations). The corresponding number of chiasmata (points of crossover) is indicated in parentheses beneath each structure.

Each configuration involves of a different number of chromosomes:

- Univalent: Involves a single chromosome, which does not pair or form any chiasmata with another chromosome.
- Rod bivalent: Involves two chromosomes, forming a linear pairing with one chiasma.
- Ring bivalent: Involves two chromosomes forming a ring-shaped pairing with two chiasmata.
- Trivalent: Involves three chromosomes, forming a more complex structure with two chiasmata.
- Rod quadrivalent: Involves four chromosomes, forming a linear configuration with three chiasmata.
- Ring quadrivalent: Involves four chromosomes forming a ring-shaped configuration with four chiasmata.

5. **Supplementary Figure 2:** Plant development under controlled conditions, highlighting a flowering delay in plant 13D compared to the others**.**

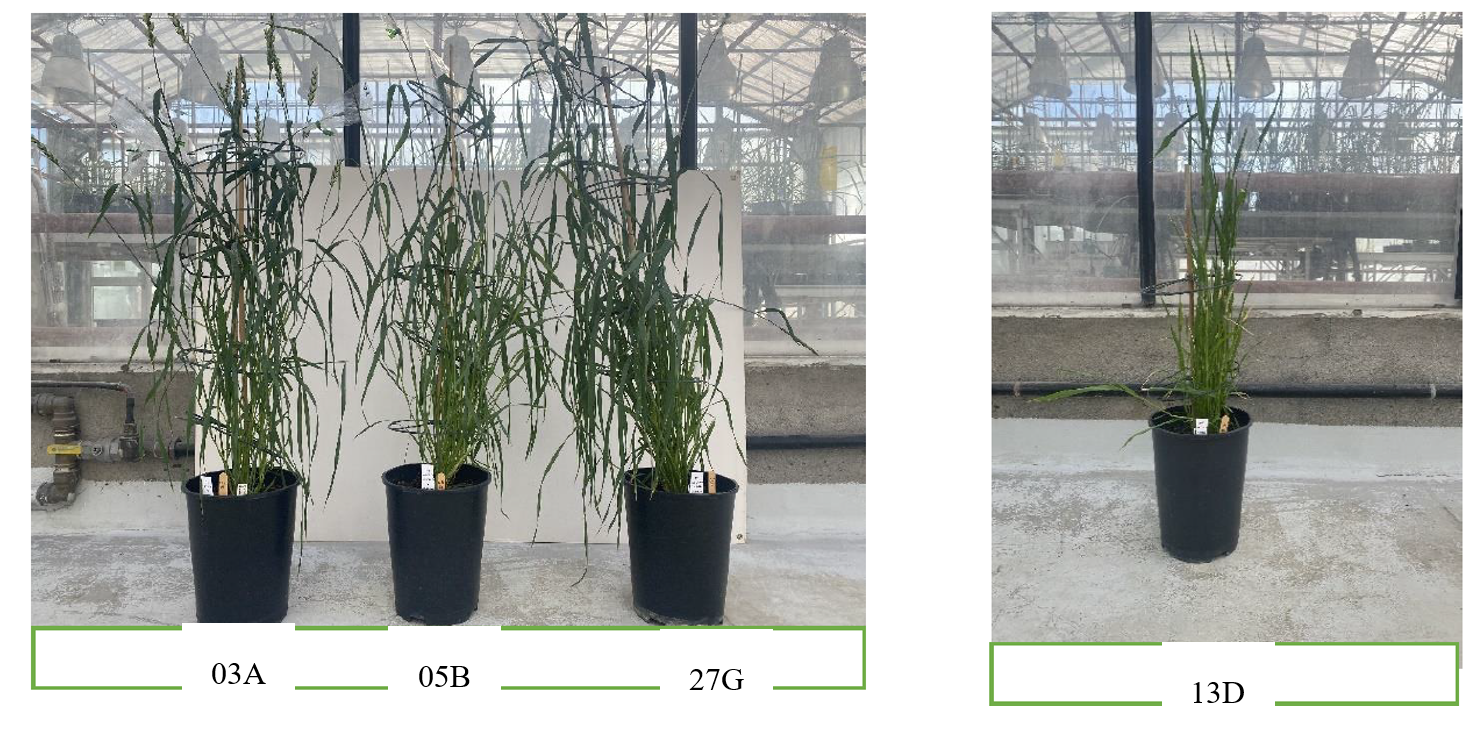

6. **Supplementary Figure 3:** Anther trio from the same flowers of individual 23F (CS ph1b/Cad msh7-3D) observed under a stereomicroscope. We observed different anther sizes within the same spikelet showing a complete desynchronization of anther development and the meiosis cycle.

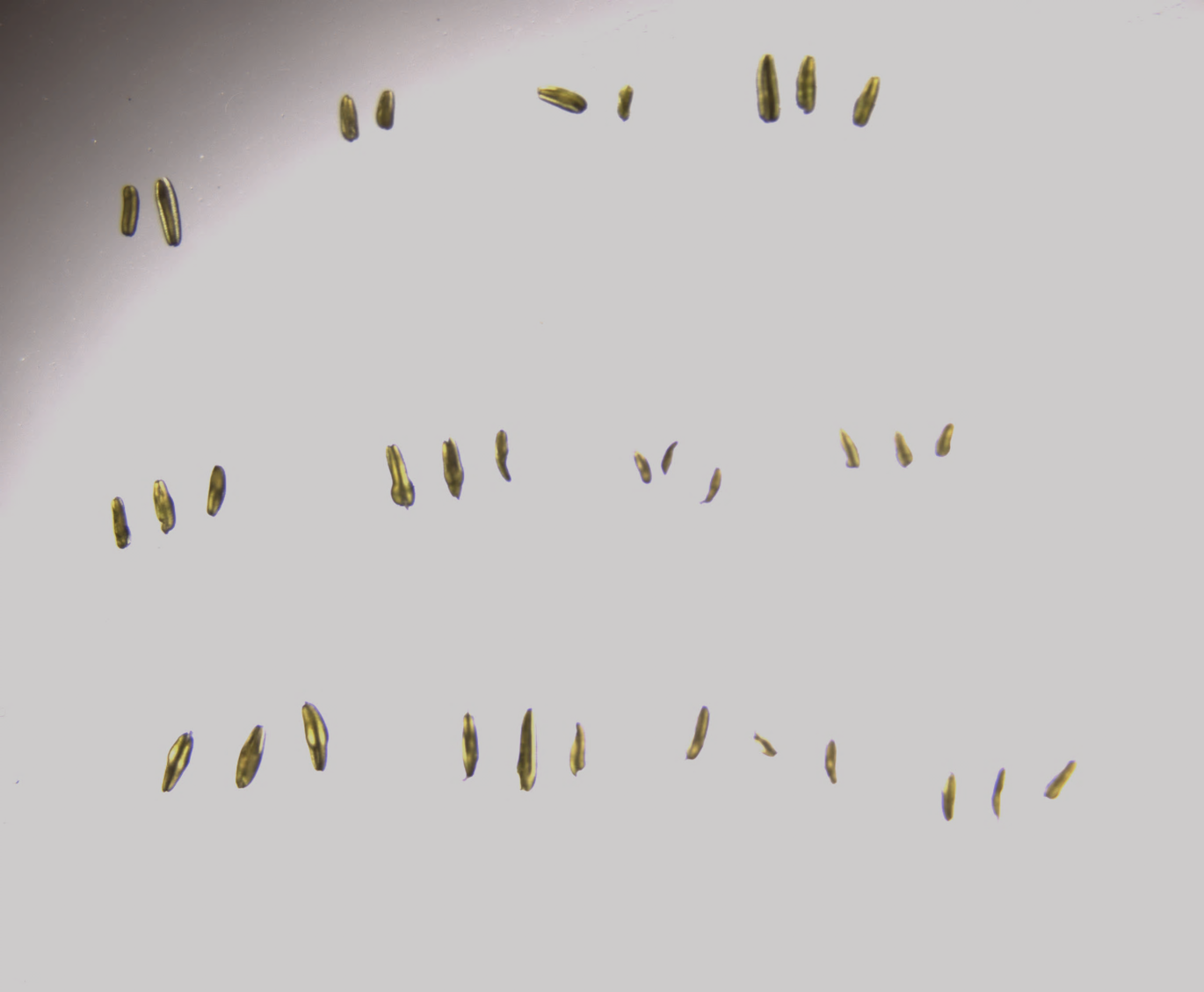

7. **Supplementary Table 4:** Matrices of Mann-Whitney Test p-values for each variable

Matrix of Mann-Whitney Test p-values for Chiasma Number: Non-significant Differences (p > 0.05) Highlighted in Yellow

|  | CS | Cad | Cadmsh7-3D | Cadzip4-5B/  CSph2b | CSph2b | Cadzip4-5B | CSph1b/  Cadmsh7-3D | Cadzip4-5B/  msh7-3D | CSph1b |
| --- | --- | --- | --- | --- | --- | --- | --- | --- | --- |
| Cad | 0.82440 | – | – | – | – | – | – | – | – |
| Cadmsh7-3D | <2e-16 | <2e-16 | – | – | – | – | – | – | – |
| Cad | <2e-16 | <2e-16 | 1.9e-07 | – | – | – | – | – | – |
| CSph2b | <2e-16 | <2e-16 | 4.0e-06 | 0.43593 | – | – | – | – | – |
| Cadzip4-5B | <2e-16 | <2e-16 | <2e-16 | <2e-16 | <2e-16 | – | – | – | – |
| CSph1b/  Cadmsh7-3D | <2e-16 | <2e-16 | <2e-16 | 2.5e-05 | 1.6e-06 | <2e-16 | – | – | – |
| Cadzip4-5B/  msh7-3D | <2e-16 | <2e-16 | <2e-16 | 3.3e-08 | 6.5e-09 | <2e-16 | 0.10093 | – | – |
| CSph1b | <2e-16 | <2e-16 | <2e-16 | 1.0e-11 | 5.3e-12 | 0.00014 | 1.6e-06 | 0.00035 | – |
| CSph1b/ph2b | <2e-16 | <2e-16 | <2e-16 | 5.2e-12 | 3.3e-12 | 0.00114 | 1.6e-07 | 6.3e-05 | 0.61010 |

Matrix of Mann-Whitney Test p-values for Univalent Number: Non-significant Differences (p > 0.05) Highlighted in Yellow

|  | CS | Cad | Cadmsh7-3D | Cadzip4-5B/  CSph2b | CSph2b | Cadzip4-5B | CSph1b/  Cadmsh7-3D | Cadzip4-5B/  msh7-3D | CSph1b |
| --- | --- | --- | --- | --- | --- | --- | --- | --- | --- |
| Cad | 0.53859 | – | – | – | – | – | – | – | – |
| Cadmsh7-3D | <2e-16 | <2e-16 | – | – | – | – | – | – | – |
| Cadzip4-5B/  CSph2b | <2e-16 | <2e-16 | 1.1e-13 | – | – | – | – | – | – |
| CSph2b | <2e-16 | <2e-16 | 9.8e-11 | 0.00184 | – | – | – | – | – |
| Cadzip4-5B | <2e-16 | <2e-16 | <2e-16 | 3.1e-09 | <2e-16 | – | – | – | – |
| CSph1b/  Cadmsh7-3D | <2e-16 | <2e-16 | <2e-16 | 0.00187 | 2.1e-11 | 0.00065 | – | – | – |
| Cadzip4-5B/  msh7-3D | <2e-16 | <2e-16 | <2e-16 | 2.1e-10 | <2e-16 | 0.17905 | 2.2e-05 | – | – |
| CSph1b | <2e-16 | <2e-16 | <2e-16 | 0.00037 | 1.1e-14 | 0.03176 | 0.28703 | 0.00292 | – |
| CSph1b/ph2b | <2e-16 | <2e-16 | <2e-16 | 7.1e-13 | <2e-16 | 2.9e-05 | 1.0e-09 | 0.00644 | 2.0e-08 |

Matrix of Mann-Whitney Test p-values for Rod Bivalent Number: Non-significant Differences (p > 0.05) Highlighted in Yellow

|  | CS | Cad | Cadmsh7-3D | Cadzip4-5B/  CSph2b | CSph2b | Cadzip4-5B | CSph1b/  Cadmsh7-3D | Cadzip4-5B/  msh7-3D | CSph1b |
| --- | --- | --- | --- | --- | --- | --- | --- | --- | --- |
| Cad | 0.0904 | – | – | – | – | – | – | – | – |
| Cadmsh7-3D | 0.0017 | 1.2e-06 | – | – | – | – | – | – | – |
| Cadzip4-5B/  CSph2b | <2e-16 | <2e-16 | <2e-16 | – | – | – | – | – | – |
| CSph2b | 3.3e-11 | 4.8e-15 | 2.0e-06 | 1.3e-07 | – | – | – | – | – |
| Cadzip4-5B | 3.2e-13 | <2e-16 | 1.3e-05 | <2e-16 | 0.1457 | – | – | – | – |
| CSph1b/  Cadmsh7-3D | <2e-16 | <2e-16 | <2e-16 | 0.1457 | 2.0e-06 | <2e-16 | – | – | – |
| Cadzip4-5B/  msh7-3D | <2e-16 | <2e-16 | <2e-16 | 0.0017 | 3.2e-13 | <2e-16 | 6.3e-07 | – | – |
| CSph1b | 3.0e-11 | 1.2e-15 | 1.2e-05 | 1.1e-07 | 0.9611 | 0.1622 | 2.1e-06 | 2.0e-14 | – |
| CSph1b/ph2b | <2e-16 | <2e-16 | 3.1e-15 | 0.1030 | 8.4e-09 | <2e-16 | 0.0018 | 0.3593 | 1.5e-08 |

Matrix of Mann-Whitney Test p-values for Ring Bivalent Number: Non-significant Differences (p > 0.05) Highlighted in Yellow

|  | CS | Cad | Cadmsh7-3D | Cadzip4-5B/  CSph2b | CSph2b | Cadzip4-5B | CSph1b/  Cadmsh7-3D | Cadzip4-5B/  msh7-3D | CSph1b |
| --- | --- | --- | --- | --- | --- | --- | --- | --- | --- |
| Cad | 0.14401 | – | – | – | – | – | – | – | – |
| Cadmsh7-3D | <2e-16 | 3.9e-16 | – | – | – | – | – | – | – |
| Cadzip4-5B/  CSph2b | <2e-16 | 3.4e-15 | 0.07207 | – | – | – | – | – | – |
| CSph2b | <2e-16 | 1.4e-14 | 0.36188 | 0.01281 | – | – | – | – | – |
| Cadzip4-5B | <2e-16 | <2e-16 | <2e-16 | <2e-16 | 1.4e-14 | – | – | – | – |
| CSph1b/  Cadmsh7-3D | <2e-16 | <2e-16 | 0.96112 | 0.01654 | 0.34392 | <2e-16 | – | – | – |
| Cadzip4-5B/  msh7-3D | <2e-16 | <2e-16 | 0.18854 | 0.37452 | 0.04135 | <2e-16 | 0.07095 | – | – |
| CSph1b | <2e-16 | <2e-16 | 7.6e-07 | 8.2e-12 | 0.00062 | 2.6e-07 | 4.4e-09 | 6.3e-12 | – |
| CSph1b/ph2b | <2e-16 | <2e-16 | 0.05034 | 0.00016 | 0.37452 | 7.7e-14 | 0.02582 | 0.00050 | 0.00448 |

Matrix of Mann-Whitney Test p-values for Complex Form Number: Non-significant Differences (p > 0.05) Highlighted in Yellow

|  | CS | Cad | Cadmsh7-3D | Cadzip4-5B/  CSph2b | CSph2b | Cadzip4-5B | CSph1b/  Cadmsh7-3D | Cadzip4-5B/  msh7-3D | CSph1b |
| --- | --- | --- | --- | --- | --- | --- | --- | --- | --- |
| Cad | – | – | – | – | – | – | – | – | – |
| Cadmsh7-3D | 0.28698 | 0.26420 | – | – | – | – | – | – | – |
| Cadzip4-5B/  CSph2b | 1.2e-07 | 2.8e-08 | 2.4e-05 | – | – | – | – | – | – |
| CSph2b | 2.0e-08 | 3.8e-09 | 7.2e-06 | 0.65945 | – | – | – | – | – |
| Cadzip4-5B | 6.7e-13 | 6.0e-14 | 2.5e-09 | 0.00959 | 0.11748 | – | – | – | – |
| CSph1b/  Cadmsh7-3D | 1.5e-15 | <2e-16 | 1.5e-11 | 1.1e-05 | 0.00187 | 0.01359 | – | – | – |
| Cadzip4-5B/  msh7-3D | <2e-16 | <2e-16 | 1.1e-12 | 4.9e-07 | 0.00023 | 0.00114 | 0.62002 | – | – |
| CSph1b | 1.5e-15 | <2e-16 | 9.7e-12 | 3.2e-06 | 0.00029 | 0.00187 | 0.37798 | 0.59804 | – |
| CSph1b/ph2b | 1.0e-13 | 8.7e-15 | 4.0e-10 | 7.0e-05 | 0.00227 | 0.01364 | 0.59649 | 0.71664 | 0.89845 |

8. **Supplementary Figure 4.** Meiotic progression during prophase I in control wheat and interspecific hybrid mutants.

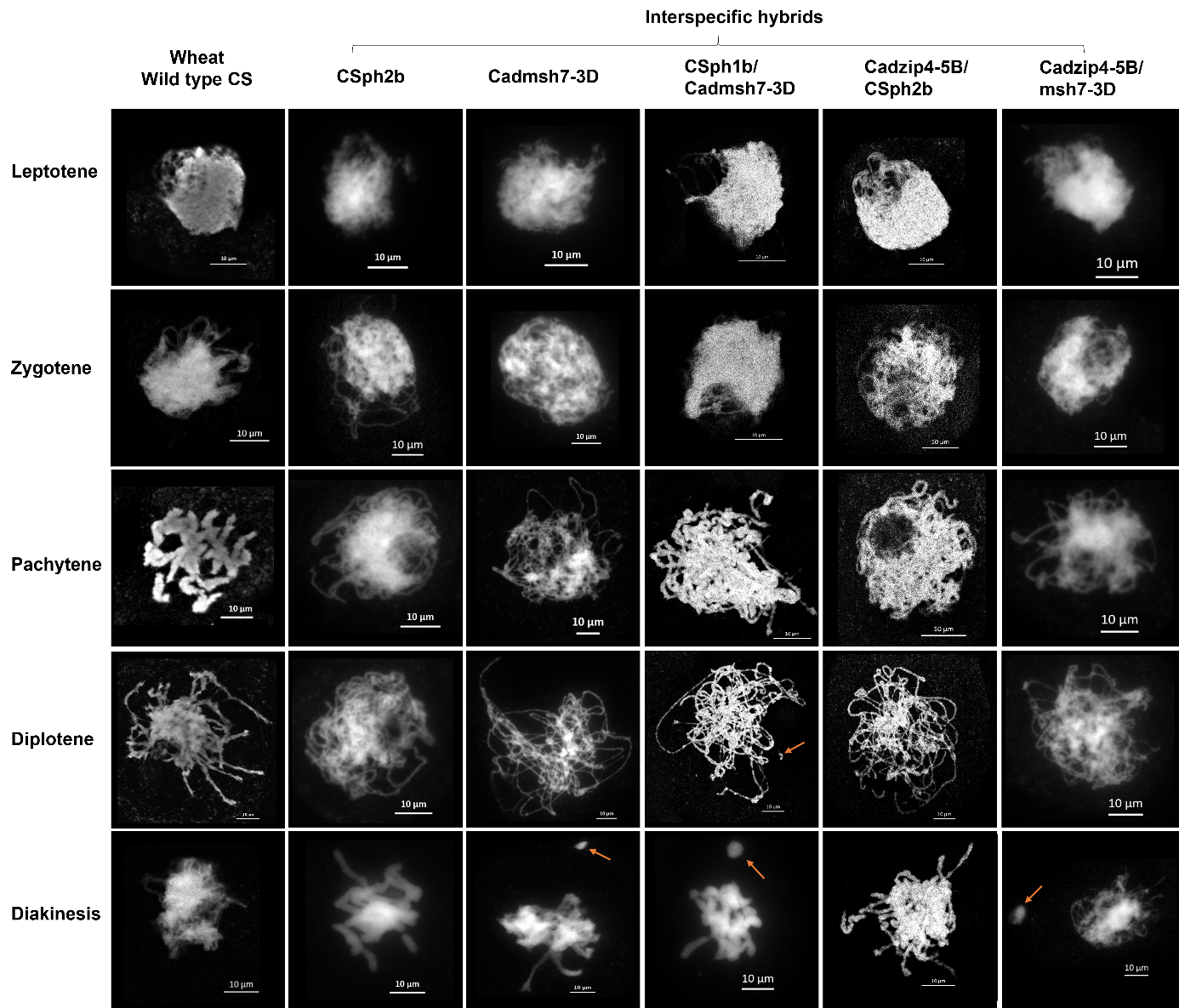

Representative images showing the sequential stages of prophase I (leptotene, zygotene, pachytene, diplotene, diakinesis) are shown following DAPI staining. Chromosome fragmentation is indicated by an arrow. Cells were observed under microscope with objective at magnification X 40, scale 10 µm.

9. **Supplementary Table 5.** Frequency of meiotic products and degree of DNA fragmentation observed in the different hybrids.

| Hybrid | Tetrads (%) | Triads (%) | Dyads (%) | Polyads (%) | Mean of fragmentation number / cell | Number of cells analyzed |
| --- | --- | --- | --- | --- | --- | --- |
| 04B  CS ph2b | 98 | 2 | 0 | 0 | 4.84 | 50 |
| 08C  Cad msh7-3D | 76.09 | 21.74 | 0 | 2.17 | 2.82 | 46 |
| 23F  CS ph1b/  Cad msh7-3D | 88 | 4 | 2 | 6 | 2.74 | 50 |
| 28G  Cad zip4-5B/  CS ph2b | 60 | 23.33 | 1.33 | 13.33 | 2.70 | 30 |
| 17E  Cad zip4-5B/  Cad msh7-3D | 80 | 10 | 6 | 4 | 4.64 | 50 |

Percentages of tetrads, triads, dyads, and polyads were calculated from the total number of post-meiotic cells analysed for each hybrid. The mean of fragmentation number was assessed from the mean number of chromosome fragments observed per cell at tetrad stage.
